# An abstract categorical decision code in dorsal premotor cortex

**DOI:** 10.1101/2022.08.24.505180

**Authors:** Gabriel Diaz-deLeon, Manuel Alvarez, Lucas Bayones, Antonio Zainos, Jerónimo Zizumbo, Sergio Parra, Sebastián Pujalte, Ranulfo Romo, Román Rossi-Pool, Victor De Lafuente

## Abstract

The dorsal premotor cortex (DPC) has classically been associated with a role in preparing and executing the physical motor variables during cognitive tasks. While recent work has provided nuanced insights into this role, here we propose that DPC also participates more actively in decision-making. We recorded neuronal activity in DPC while two trained monkeys performed a vibrotactile categorization task, utilizing two distinct ranges of stimuli values that varied on two physical attributes: vibrotactile frequency and amplitude. We observed a broad heterogeneity across DPC neurons, the majority of which maintained the same response patterns across attributes and ranges, coding in the same periods, mixing temporal and categorical dynamics. The predominant categorical signal was maintained throughout the delay, movement periods and notably during the inter-trial period. Putting the entire population’s data through two dimensionality reduction techniques, we found that imposing the sensory structure yielded pure categorical and temporal representations. Furthermore, projecting the activity of one population over the population axes of the other yielded identical categorical and temporal responses. Finally, we sought to identify functional subpopulations based on the combined activity of all stimuli, neurons, and time points, however we found a continuum of single-unit responses mixing temporal and categorical dynamics. All this points to DPC playing a more decision-related role than previously anticipated.

**SIGNIFICANCE STATEMENT:** The DPC’s role in the somatosensory processing network has been generally limited to movement, but our current results suggest a more abstract function. We recorded DPC’s activity in two monkeys trained in a vibrotactile categorization task of two distinct physical attributes, and found a strong decision signal throughout the population, underpinned by purely temporal signals. Importantly, this abstract decision signal remains during the inter-trial period suggesting a consolidation role. Neurons maintained consistent and significant responses for both attributes, and the entire population activity converged to identical categorical representations, even when cross-projected between two contexts. These results suggest that DPC plays a larger role during decision-making and consolidation, regardless of the stimulus attributes that triggered the decision report.

## INTRODUCTION

Daily, humans and animals learn to recognize commonalities between distinct subjects and produce unifying categories and concepts in a natural fashion, instead of storing every individual experience. This process, defined as categorization, permits the approximation of quick decisions necessary in daily life, e.g., labeling the worth of an item as expensive or slowing down when we realize we are driving too fast in a school zone. Once learned, these categories can be generalized, assimilating unrelated groups, which results in the expansion of the previous category’s definition. Thus, the understanding of novel topics and experiences, and matching them with the most appropriate decisions, depends on prior knowledge of the category in question. Furthermore, this categorization process also depends on the contextual situation; under different conditions, the same category could have different meanings. For example, acceptable driving speeds depend on context, given that they differ between a city street and a highway. The interval values that delimit each category are modulated in a context-dependent manner.

Complex phenomena such as categorization and decision-making rely on a variety of underlying mechanisms. Although it is known that specific processes depend on the decision to be made (1–3), it is believed that some common task-resolving mechanisms could emerge as well (4, 5). Neural coding from frontal lobe areas demonstrates a high capacity for abstraction and generalization (4, 6–9), which may even be generalized across different sensory modalities (10). In contrast, sensory cortices respond faithfully or code specific attributes of the stimuli (11). A previous study in individual neurons in the primary somatosensory cortex (S1), in which trained monkeys had to categorize three different sensory features of a vibrotactile stimulus (VCT), showed that some neurons code a particular feature of the stimuli while others code all features indiscriminately (12). Thus, S1 neurons represent different aspects of the stimuli, but not its category.

Employing the same categorization task (VCT), here, we studied the neuronal activity recorded from the dorsal premotor cortex (DPC). This cortical region was historically associated with the planning of physical movement and response (13–20). Several studies suggest that the activity in DPC can also be related to an abstracted representation of the final decision, making this area a prime candidate to study the categorical representation that emerges within the cortex. In particular, during a temporal pattern discrimination task, neurons of this area codify stimulus patterns in abstracted categories, which were signaled in several different epochs of the task, such as the working memory, comparison, and decision-making periods (4). Moreover, a variety of heterogeneous activity with a broad diversity and mixed selectivity dynamics is elicited (21). To give sense to this diversity, the neural dynamics were investigated at the population level, employing the collective activity of the DPC network during the decision process (22). This population level approach was also used to analyze DPC responses during a visual task, showing that preparatory and movement dynamics were coded in orthogonal population dynamics (20, 23). Importantly, at different times, the same neurons modify their responses to generate these separate coding subspaces (20). These recent findings provide relevant evidence to revindicate the role of DPC during abstract categorization, working memory and decision making. Therefore, the participation of DPC in more cognitive processes should be further explored.

In this regard and since it is known that categories serve as a mechanism for uniting various dissimilar objects, we posed two initial questions to face this exploration: is it possible that separate members of the same category are represented similarly in the neuronal activity? Does the brain unite separate categorical representations, or maintain them independently in isolated subpopulations? To address these questions, we trained two monkeys to perform the VCT, which consisted in evaluating the magnitude of a given stimulus as “high” or “low” based on the physical attribute that was varied (frequency or amplitude). For each attribute, two different superimposed ranges were trained. Importantly, stimulus magnitudes shared by sets of the same physical attribute should be categorized as “low” in one while “high” in the other.

Characterizing the breadth of responses that appear among the population’s single neurons, we observed an overwhelming categorical representation that was maintained throughout the delay period, lasting until the inter-trial period. Further, all observable sensory dynamics appeared to be correlates of the categorical responses, consistently across the 2 physical attributes and the two superimposed sets. Then, we transit to considering how a common population of single units behave for differing stimulus attributes. This would allow us to see if neurons were committed to a single attribute, or if they would present with a more abstract response pattern that unifies the distinct physical attributes. We found that most neurons presented similar response patterns for both attributes, often coding for both attributes at the same moment.

Although single-unit studies still yield interesting findings in this area, little is known about the aggregate activity that emerges across the entire population when the VCT is performed. To address this issue we used dimensionality reduction techniques to better understand the heterogeneity of dynamics occurring within the population recorded from DPC, and find the most representative underlying dynamics (20, 22, 24–26). Once again, we observed a consistently maintained categorical representation and the dynamics that emerge from different sets are entirely relatable using populations of neurons recorded between different sets. Moreover, two orthogonal dynamics were observed, one during the delay and the other one during the movement/inter-trial periods. Despite this, we found that these orthogonal population responses are represented by the same neural substrate; instead of clusters of neurons associated with each orthogonal dynamic, we find a continual transition between these signals. These results demonstrate that DPC wholly leverages its heterogeneous activity to form a highly efficient abstract decision categorical code, which serves to categorize different stimulus attributes regardless of their physical attributes.

## RESULTS

### Vibrotactile Categorization Task

Two monkeys *(Macaca mulatta)* were trained to categorize a vibrotactile stimulus as high or low for two different physical attributes: frequency and amplitude, each in two different overlapping ranges: short frequency range (SFR= 10-30 Hz in steps of 2 Hz) and short amplitude range (SAR = 20-80 μm in steps of 6 μm); long frequency range (LFR= 14-78 Hz in steps of 8 Hz) and long amplitude range (LAR = 42-136 μm in steps of 12 μm), resulting in a total of 4 different predetermined sets (Fig. 1A, C) (12). Sets were presented in a random order (see Methods). Monkeys’ performance was consistent for all sets: frequency short (SFR, n=61 sessions, 90.3±0.16%) and long (LFR, n=16 sessions, 82.9±1.2%, Fig. 1B); amplitude short (SAR, n=48 sessions, 85.4±0.35%) and long (LAR, n=11 sessions, 81.1±1.41%, Fig. 1D). This is represented by the rightward shift of the lighter colored psychometric curve as well as the saturation at values of 0 and 1. All data (355 neurons) was recorded from DPC (Fig. 1E), where 275 neurons were recorded during the SFR set, 232 neurons during the SAR set, 84 during the LFR set and 59 during the LAR set (Fig. 1F).

**Figure 1.**
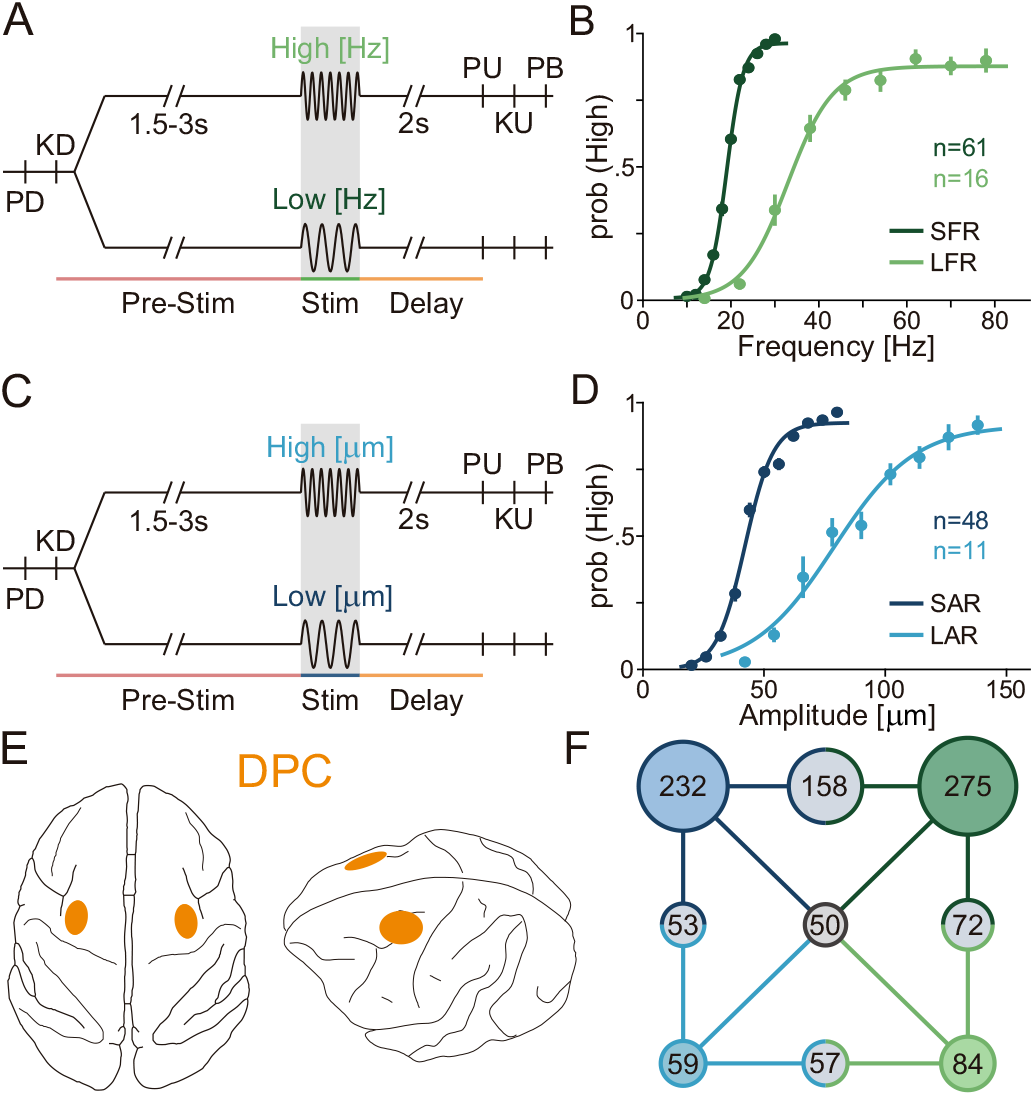
Task schematic, Psychophysical Performance, Data Sets and Recording Sites. (A, C) Task schematic for categorization of frequencies (A) and amplitudes (C). Trials start when the mechanical probe lowers, indenting the glabrous skin of one fingertip of the monkeys’ restrained right hand (probe down event, “PD”). The animals respond by placing their left hand on an immovable key (key down event, “KD”). After KD, a variable period (1.5-3s) is presented, followed by the presentation of a single stimulus lasting 0.5s. After stimulation, a fixed delay period of 2s is presented, followed by the probe up event (“PU”). PU serves as the “go” cue for monkeys to remove its hand from the key (key up event, “KU”) and report their decision using one of the two push-buttons placed in front of him (push-button event, “PB”). Correct answers were rewarded with a few drops of juice. (B, D) Psychometric performance for all categorization sets: (B) Psychometric curves for the two frequency-range sets, short (dark green) and long (light green). The short frequency set (SFR) had 12 stimuli that varied between 10 and 30 Hz, in steps of 2Hz, with the middle stimulus (20Hz) repeated once for each category (61 sessions). The long frequency range set (LFR) had 10 stimuli that varied between 14 and 78 Hz, in steps of 8Hz, with the middle stimulus (46Hz) repeated once for each category (16 sessions). (D) Psychometric curves for two amplitude-range sets, short (dark blue; SAR) and long (light blue; LAR). The SAR set had 12 stimuli that varied between 20 and 80μm, in steps of 6μm, with the middle stimulus (50μm) repeated once for each category (48 sessions). The LAR set had 10 stimuli that varied between 42 and 138μm, in steps of 12μm, with the middle stimulus (90μm) repeated once for each category (11 sessions). (E) Top (left) and lateral (right) view of a monkey brain, with recorded DPC area highlighted (yellow). (F) Diagram of populations of neurons recorded per set. Colors are consistent with those in panels B and D. Corner circles with solid colors represent sets, intersecting circles with 2 mixed colors represent populations recorded in both sets, and the black circle in the middle is the population of neurons recorded in all 4 sets.

### Single Neuron Responses during the Vibrotactile Categorization Task

Previous studies in DPC have shown that its neural activity is heterogeneous, spanning from purely categorical to purely temporal (20, 22, 27). To study this diversity, we characterized the coding and response properties of each neuron over the course of the VCT. Exemplary neurons are shown in Figs 2 and S1. We can observe a neuron with a purely categorical persistent response to the SFR set in Fig. 2A (top); it was silent for high stimulation frequencies but maintained the response during low ones. It is important to notice, in the raster plot and rate profile (Fig. 2A, bottom), that the categorical response emerged at the end of the stimulation period and was maintained throughout the delay. Remarkably, this neuron had a nearly identical response to amplitude categorization (SAR set, Fig. 2B). This phenomenon was consistently observed across neurons; in general, most had the same responses regardless of the physical attribute tested.

**Figure 2.**
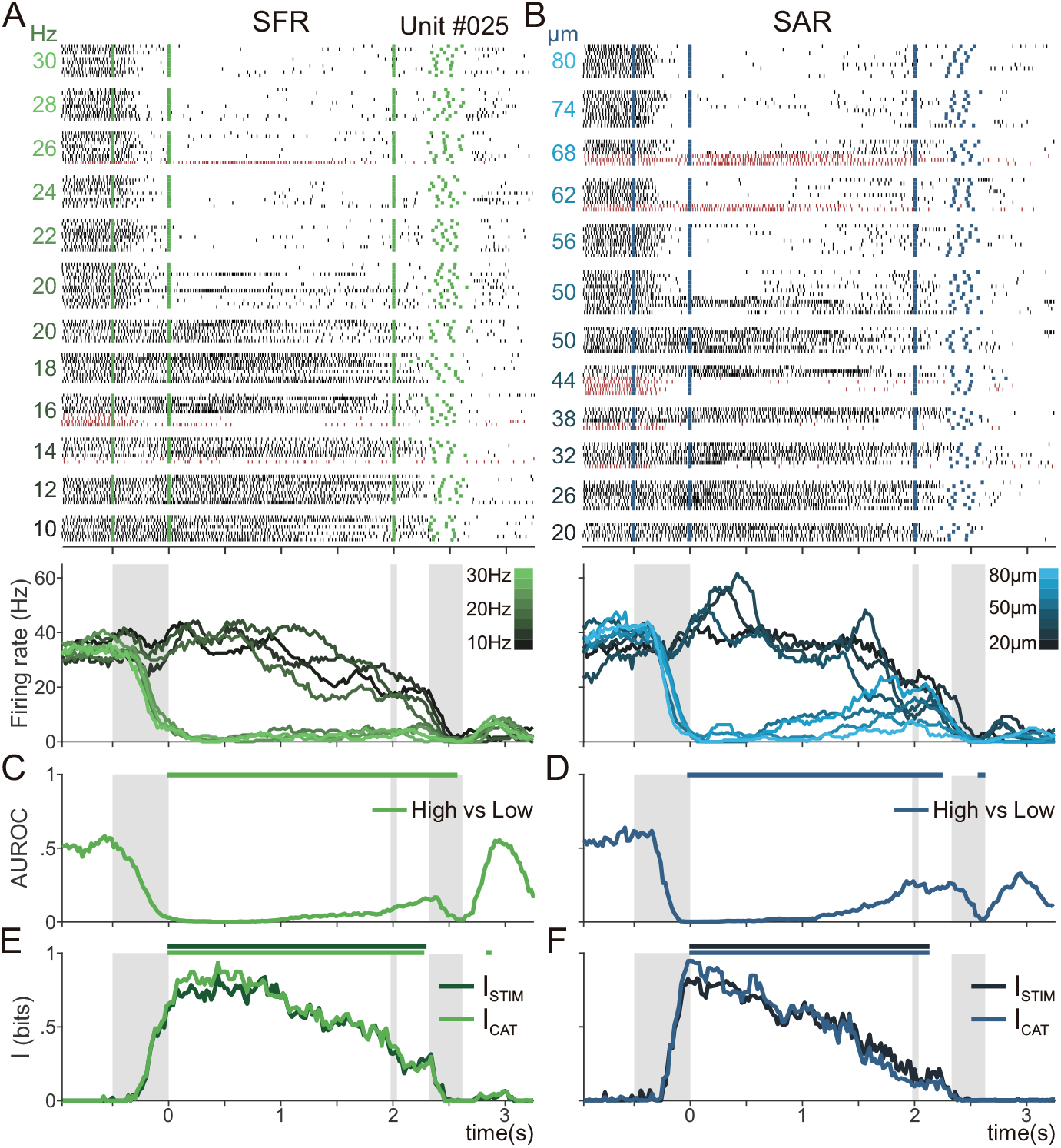
A Single Neuron’s Activity during Frequency and Amplitude categorization. (A, B) Raster plots of a single neuron recorded in the short frequency range (SFR; A, C, E) and short amplitude range (SAR; B, D, F) sets. Black and red tick marks represent spikes in trials with correct and incorrect answers, respectively. Green (A) and blue (B) tick marks indicate psychophysical events. Trials are grouped by class (stimulus intensity), with intensity values marked on the left side. From left to right, the first green and blue tick marks that occur at −0.5s indicate stimulus onset, and the second marks at t=0s indicate stimulus offset. After a fixed 2s delay period, the third green and blue tick marks indicate PU, while the 4^th^ represents KU and the 5^th^ represents PB. Figures below raster plots represent firing rate averages for all hit trials associated with each of the 8 visualized classes. Some classes were excluded for clarity in the visualization. Large grey rectangles represent the stimulation period, while thin rectangles mark PU, and the third and last rectangle marks the average movement period, beginning with KU and ending with PB. Firing rate curves range from dark grey for low category stimuli to bright green (A, frequency) or bright blue (B, amplitude) for high category stimuli. (C, D) Time-dependent differential AUROC (High vs Low) values were taken between the firing rate distributions of correct trials. AUROC<0.5 or AUROC>0.5 indicates an increased firing rate response for classes of category Low or High, respectively. Significant windows, based on permutation tests and correcting for multiple comparisons, are marked above the curve (p<0.01). (E, F) Time-dependent category (I_CAT_, light green and blue) and stimulus mutual information (I_STIM_, dark green and blue) curves (p<0.05).

To quantify the categorical nature of neuron responses we computed the AUROC (See Methods) at each time (28), between the firing rate distributions corresponding to high and low stimulus categories. For some neurons (see Fig. 2C and 2D), categorical representation during the SFR and SAR sets significantly emerges at the end of the stimulation period and persists throughout the delay until after the PU event. Its activity represents the high stimulus categories, characterized by an AUROC<0.5.

Given that a categorical code implies a transformation from the sensory representation, we wondered if pure sensory information is still present in DPC neurons. To elucidate this question, we computed the category (I_CAT_) and stimulus (I_STIM_) mutual information for each physical attribute, for each single neuron (See Methods). Surprisingly, these two calculations yielded identical results (Fig. 2E and 2F) for their values as well as their significant time windows (p<0.05). We can surmise that I_STIM_ is highlighting the same differences between the stimuli as I_CAT_ is between the categories. Moreover, this is also found for the long stimulus sets (SFR and LAR), where we can again observe that this neuron preferentially responds to low classes for both physical attributes (Fig. S1A and S1B). These results indicate that this exemplary neuron maintains a significant categorical representation, regardless of the physical attribute or stimulation range studied.

Other DPC neurons recorded in the SFR condition also exhibited a broad repertoire of dynamics that vary over the task period (Fig. S2). For example, panel A shows a neuron that responds after the go-cue, with slightly categorical dynamics in the movement period that become even more pronounced in the inter-trial period. However, in B there is a neuron whose response is slightly categorical during the stimulation period but whose clearest categorical response occurs only after movement, during the inter-trial period. The neuron in C has pure temporal dynamics up until the go-cue, becoming categorical during the movement period, and then presents mixed coding in the inter-trial period. The neuron in panel D mirrors the one shown in Fig. 2 and S1, with a persistent categorical code throughout the delay period but with a preference for the low rather than high category. Interestingly, its categorical coding is quenched after the go-cue and movement period. The neuron in Fig. S2E demonstrates a slight categorical response during the delay and after the movement period, with clear temporal modulation. Finally, panel F shows a neuron that represents purely temporal dynamics, with almost no variation between the classes throughout the entire task period. It is important to note that following the movement period, dynamics continue to emerge in several neurons (Fig. S2B, S2E, see Fig S5B [#085, #154, #169, #150, #143, #127]) in a manner that is independent of those present during the motor report. This signal is maintained even later than the reward delivery (~40ms after push button), during the inter-trial period. In summary, as opposed to (12), DPC neurons display a diverse range of categorical and temporal dynamics in their responses.

### Single Neuron Coding Dynamics

Since we now have a picture of single neuron responses in DPC, we proceed to analyze how the entire population behaves as a function of time. For each metric (AUROC, I_CAT_ and I_STIM_) and stimulus set, Fig. 3 shows the proportion of all neurons yielding significant results across the entire population. Notably, in Fig. 3A for SFR, the response pattern is similar across all 3 metrics: the categorical representation emerges at the end of the stimulation period, it is maintained persistently throughout the delay, and then the representation is reinforced during the movement period. Intriguingly, the greatest proportion of neurons with significant coding is found after movement, in the inter-trial period. The peak during the stimulus period corresponds to the emergence of a categorical decision, where the subject decides its report, and the persistent dynamic during the delay corresponds to the subject’s maintenance of the decision in working memory. The re-emergence of the categorical representation after the go-cue is in part related to the physical report itself (23). However, a high proportion of this categorical coding persists into the inter-trial period. This was previously found in DPC during a different task (4, 22), and is probably related to learning or reward processes, such as adapting the evaluation of choice outcomes into future decisions (29, 30). Note that this same response pattern was observed for the population recorded during the LFR condition (Fig. 3B).

**Figure 3.**
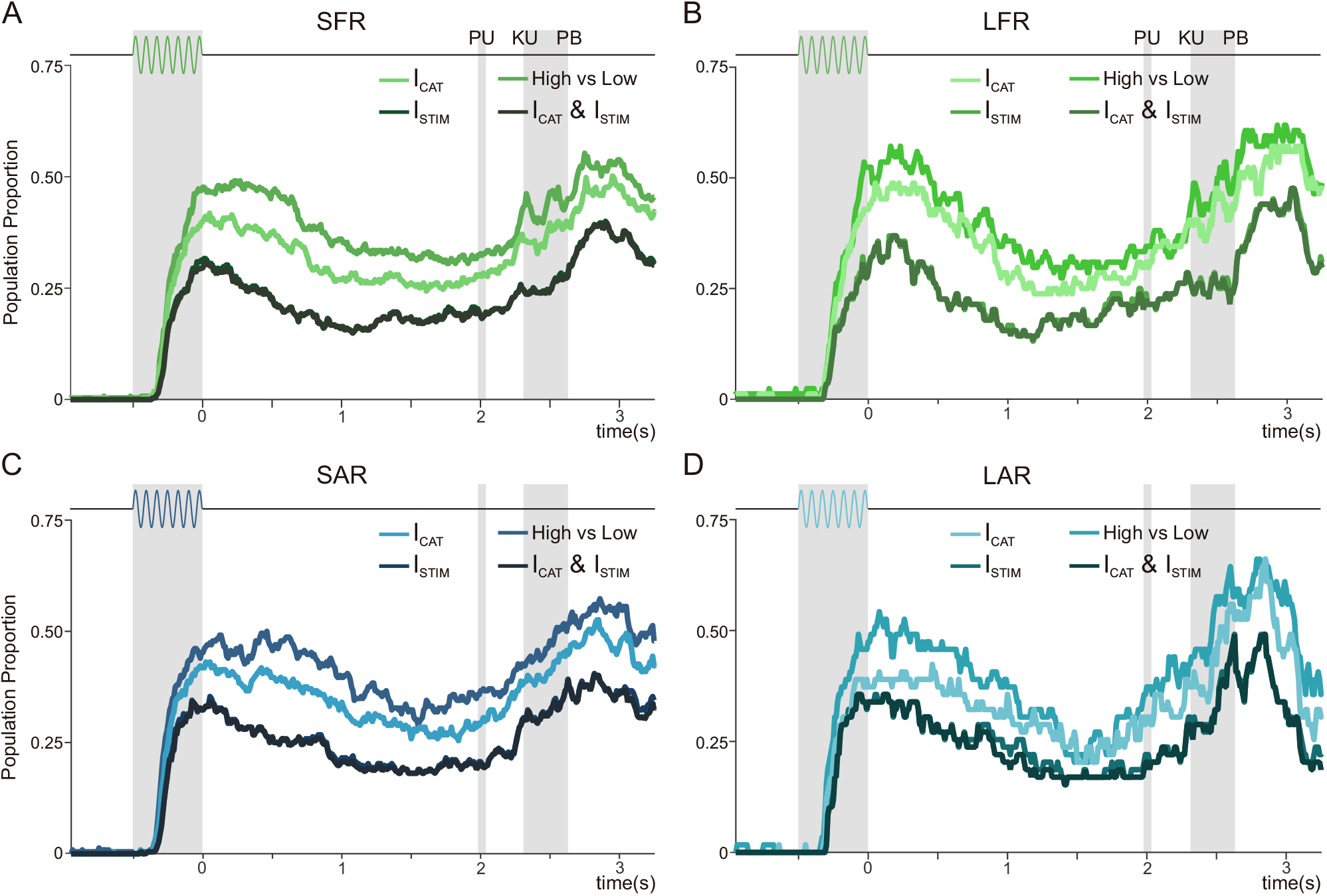
Single Neuron Coding during Frequency and Amplitude Categorization. (A-B) Neuron population coding for the stimulus frequency (A-B) and stimulus amplitude sets (C-D). Population proportions recorded for frequency short-range (SFR; A, n=275) and long-range (LFR; B, n=84) and for amplitude short-range (SAR; C, n=232) and long-range (LAR; D, n=54) with significant category information (I_CAT_), stimulus information (I_STIM_), both information types at once (I_CAT_ and I_STIM_), and differential AUROC (High vs Low) (n=275).

We now wonder if the categorical (I_CAT_) and the stimulus information (I_STIM_) can account for separate underlying dynamics. In all panels of Fig. 3, the proportion of neurons with significant I_CAT_ is greater than I_STIM_, suggesting that categorical coding is more abundant. However, the percentage of neurons with both significant I_CAT_ and I_STIM_ (I_CAT_ and I_STIM_), almost perfectly overlaps with the curve for only I_STIM_. As previously discussed for individual neurons (see Fig. 2A), this result suggests that categorical information is the only one that remains in the DPC network. In other words, information related with the identity of the stimulus is no longer present in DPC. These findings remained true for all sets, regardless of attribute or range, exhibiting a general dynamic for this cortical area. However, it remains unclear whether a single neuron performs common operations between different sets, or if the network assigns specific roles to individual neurons based on the set being tested.

To test this, we analyzed the neurons recorded for more than one set in order to understand how the DPC network codes the information during the categorization of different stimulus attributes. When comparing SFR and SAR, we calculated the information tuning as a function of time for both stimulus sets, and then compared each time bin to determine if the single neurons were coding information related to a single or both stimulus attributes. Fig. 4A shows the coding as a function of time for each neuron recorded in both sets (SFR/SAR, n=158). We observed that most neurons code both attributes at some point throughout the task. To further elaborate, we computed the population proportions with significant coding as a function of time for individual or both attributes (Fig. 4C). Note, that during the task, the percentage of neurons that codes both features (I_both,CAT_) is much higher than those with a single-feature coding (I_freq,CAT_ or I_amp,CAT_). These results strengthen our hypothesis of the network’s convergent mechanism for stimulus categorization. Further, a similar abstract population coding was observed for the neurons recorded during both long ranges of stimulus sets (Fig. 4B and D, LFR/LAR, n=56). These results indicate that the network reused the coding dynamics of most of the individual units, demonstrating a unique representation of otherwise divergent categorical identities, regardless of whether the stimulus varied based on amplitude or frequency. However, we wondered whether this occurs when considering the emerging activity from the entire population of neurons regardless each stimulus feature.

**Figure 4.**
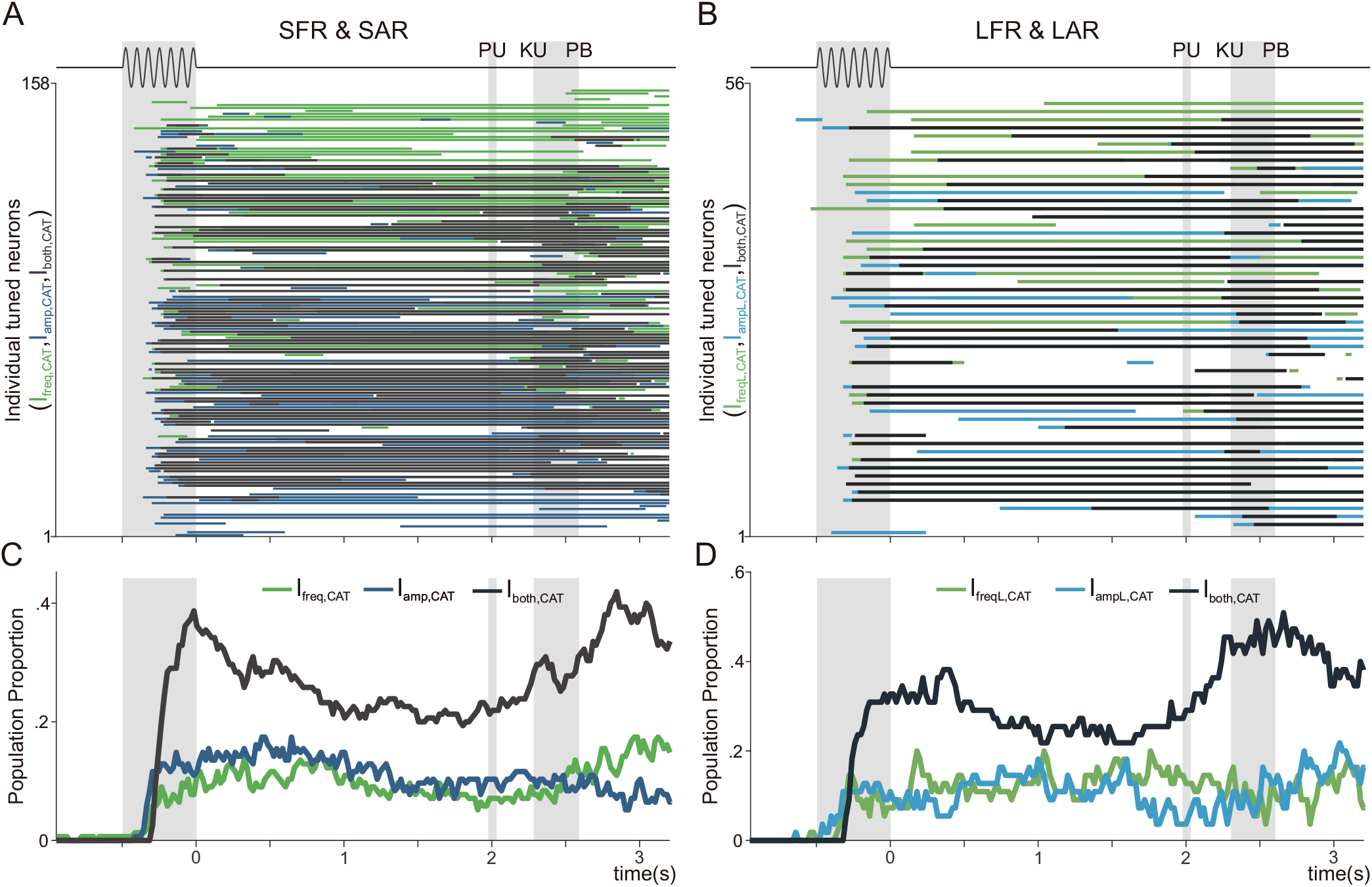
Abstract Information Coding of Stimulus Attributes. (A, B) Information tuning for all neurons recorded in the frequency and amplitude sets. (A) Neurons recorded (n = 158) during the short frequency range (SFR) and short amplitude range (SAR). (B) Neurons recorded for both long stimulus attribute ranges (LFR and LAR; n=54). Lines indicate individual neurons with significant information during only one set (blue and green) or both sets (grey). (C-D) Percentage of neurons, recorded during short-range (C) or long-range sets (D), with only frequency (green), amplitude (blue), or dual information coding (dark grey).

### Generic Population Dynamics across stimulus Attributes

To understand the signals that were obfuscated in the heterogeneity of single neuron responses, dimensionality reduction methods were recently applied to compute the relevant population dynamics (25, 31–33). Here, we employed demixed-Principal Component Analysis (dPCA) to find the population axes that maximize the amount of explained variance associated with a particular task parameter (22, 24). To compute these representative axes (dPCs), we calculated the marginalized covariance matrices which summarize the population coactivity with respect to each task parameter. These decoding axes yield dimensions that summarize the population-level dynamics, allowing us to understand how the network is representing the different parameters associated with the task. Moreover, this approach allowed us to separate the population coding dynamics from the pure temporal signals that explained most of the variance.

We begin by considering the four stimulus sets separately (SFR, SAR, LFR, and LAR) and computing the dPCs with the marginalized activity with respect to the stimulus identity. In Fig. 5 panels A and B (top and bottom), we plotted the first two stimulus-dPCs calculated for SFR and SAR, where each one has 8 classes projected. In a similar manner to the single neuron analysis, the stimulus-axes for SFR and SAR showed a categorical representation that emerges halfway through the stimulation period and maintain itself throughout the delay period until after the GO cue. It is only during the movement period that the dPCs began to diverge: the 1^st^ dPC maintains its categorical representation until well-after the GO cue and movement period (Fig. 5A and 5B, top), while the 2^nd^ dPC exhibits its strongest categorical representation during the inter-trial period (Fig. 5A and 5B, bottom). Remarkably, this same response pattern was observed for the LFR and LAR sets (Fig. S5A and B, top and bottom). Since each dPCs was identified using separate data sets, the congruence across these results strongly suggests that a common, generic coding mechanism was being employed.

**Figure 5.**
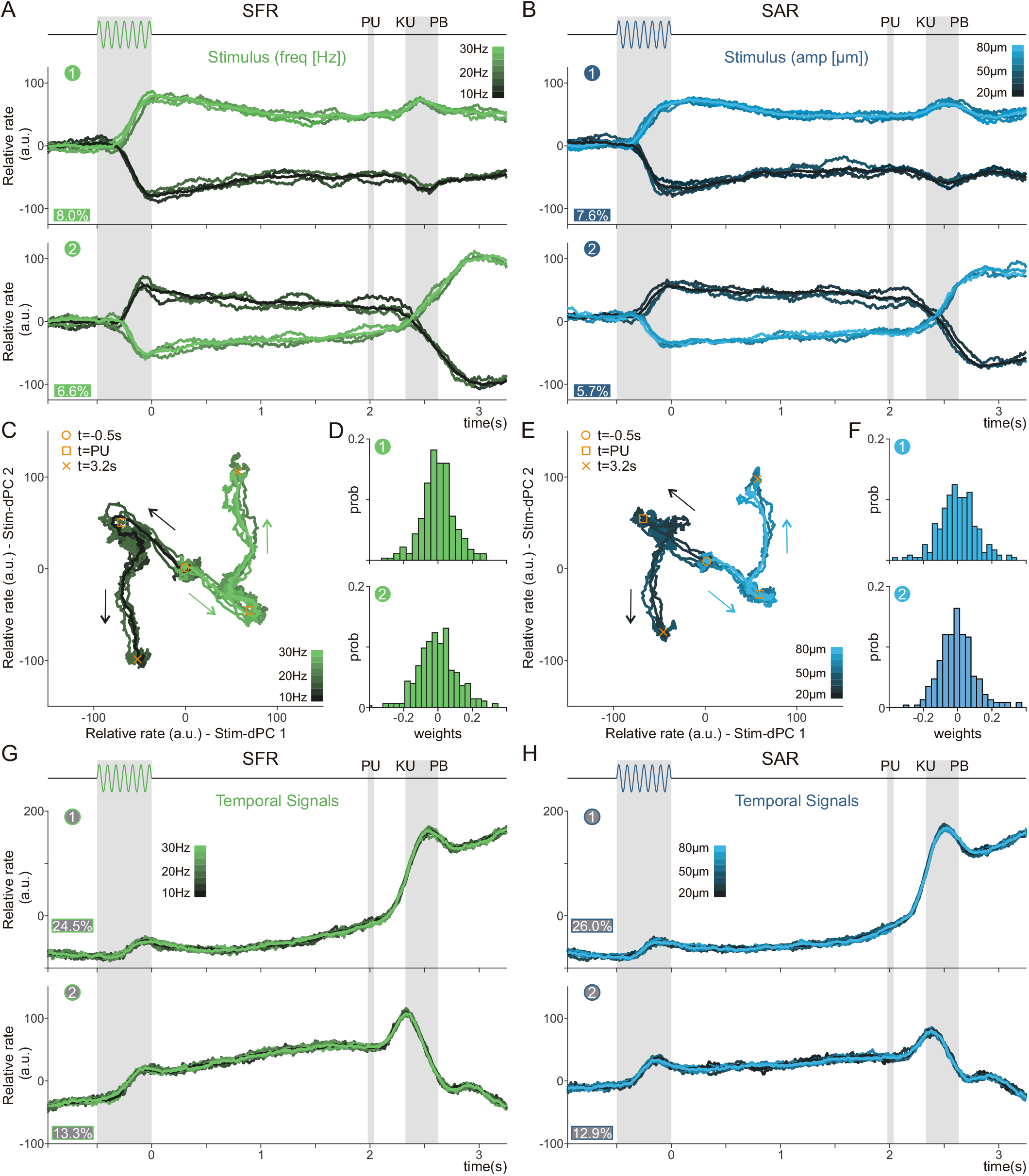
Stimulus and Temporal Population Dynamics. (A, B) Projections of class averages over the first and second decoding stimulus-axes of the short-range population for frequency (A, SFR, n=275) and amplitude (B, SAR, n=232). Classes ranging from dark (lowest stimulus) to light (highest stimulus). The ordinal number of each dPC is shown in a circle; explained variances are shown as percentages. (C, E) Phase diagram of the same classes projected across the first 2 decoding axes for frequency (C) and amplitude (E). The yellow circle marks t=-0.5s (SO), the yellow square marks the end of the delay (PU, t=2s), and the yellow x marks t=3.2s (inter-trial period). (D, F) Distributions of the neuronal weights for the first 2 decoding axes of short-range for frequency (SFR, D) and amplitude (SAR, F). (G, H) The first two temporal population signals for the short-range population for frequency (SFR, G) and amplitude (SAR, H).

If we consider both stimulus-dPCs together, it is possible to understand how the network manages to differentiate the traces related to each category during the different task periods. In panels C and E of Fig. 5, we plot two-dimensional phase diagrams obtained from the first two dPCs associated with frequency and amplitude, respectively. All traces start together in the center of the phase diagram, but soon afterwards the traces for the stimulus related to category low and high begin to diverge, consolidating into what could be separate decision paths. Importantly, after the PU event (square symbol), the population evolution changed drastically. This suggests that the population dynamics between these two task periods are almost orthogonal (20, 23, 34). Intriguingly, this orthogonal population signal endures well into the inter trial period. Moreover, we also found that this same behavior remains consistent for the long-range sets as well (Fig. S3C and E for LFR and LAR, respectively).

Considering the contributions of each neuron for these two dPC axes, we find that the weights are normally distributed around 0, indicating that the network balances positive and negative coding equally (Fig. 5D and F and Fig. S3D and F). We then computed the pure temporal population responses, employing the condition-independent marginalized matrices to calculate the first two temp-dPCs (24). For all sets, these two signals remained unaltered and explained more than the 42% of the whole variance (Fig. 5G and H and Fig. S3G and H). This high percentage suggests that they constitute an essential underlying mechanism for this categorization process (27). When considered together, these results suggest that the same coding and temporal population dynamics are exploited by DPC neurons during the categorization of different stimulus attributes and ranges.

Finally, we decided to go further and test the universality of these abstracted population codes. For this, we produced the decoding axes for a common population of neurons recorded in both the SFR and SAR sets and projected then the SAR population activity over the SFR decoding axes (Fig. 6A), and vice versa (Fig. 6B). The axes (dPC) that were optimized to decode stimulus identity during the SFR set, was employed to project the population activity recorded during the SAR set. This methodological procedure is only possible because the neurons were recorded during both sets: SFR and SAR. We also performed the same procedure for the common population between the LAR and LFR sets (Fig. S4A and B). The projection of SAR-population activity over the 1^st^ and 2^nd^ SFR-decoding axes gives rise to the same categorical representations as in Fig. 5: the categorical decision coding last through the delay and grows in the inter-trial period (Fig. 6C). This remains consistent when projecting the LAR-population activity over the LFR-decoding axes (Fig. S4C). Moreover, a nearly identical categorical representation appears when we project the SFR-population activity over the SAR-decoding axes (Fig. 6D), as well as when we project the LFR-population activity over the LAR-decoding axes (Fig. S4D). Furthermore, the temporal signals for the SAR-population activity projected over SFR-temporal axes (Fig. 6E) mirror the SFR-population activity projected over the SAR-temporal decoding axes (Fig. 6F). Analogous results were also found for long range sets (Fig. S4E and F). Moreover, these signals are also remarkably similar to the pure decoding axes that were obtained for the entire populations in Fig. 5 and Fig. S3. These results strengthen the evidence that the DPC network uses a universal population dynamic during this task. The same coding and temporal population response are triggered regardless of the stimulus’ physical attribute or range.

**Figure 6.**
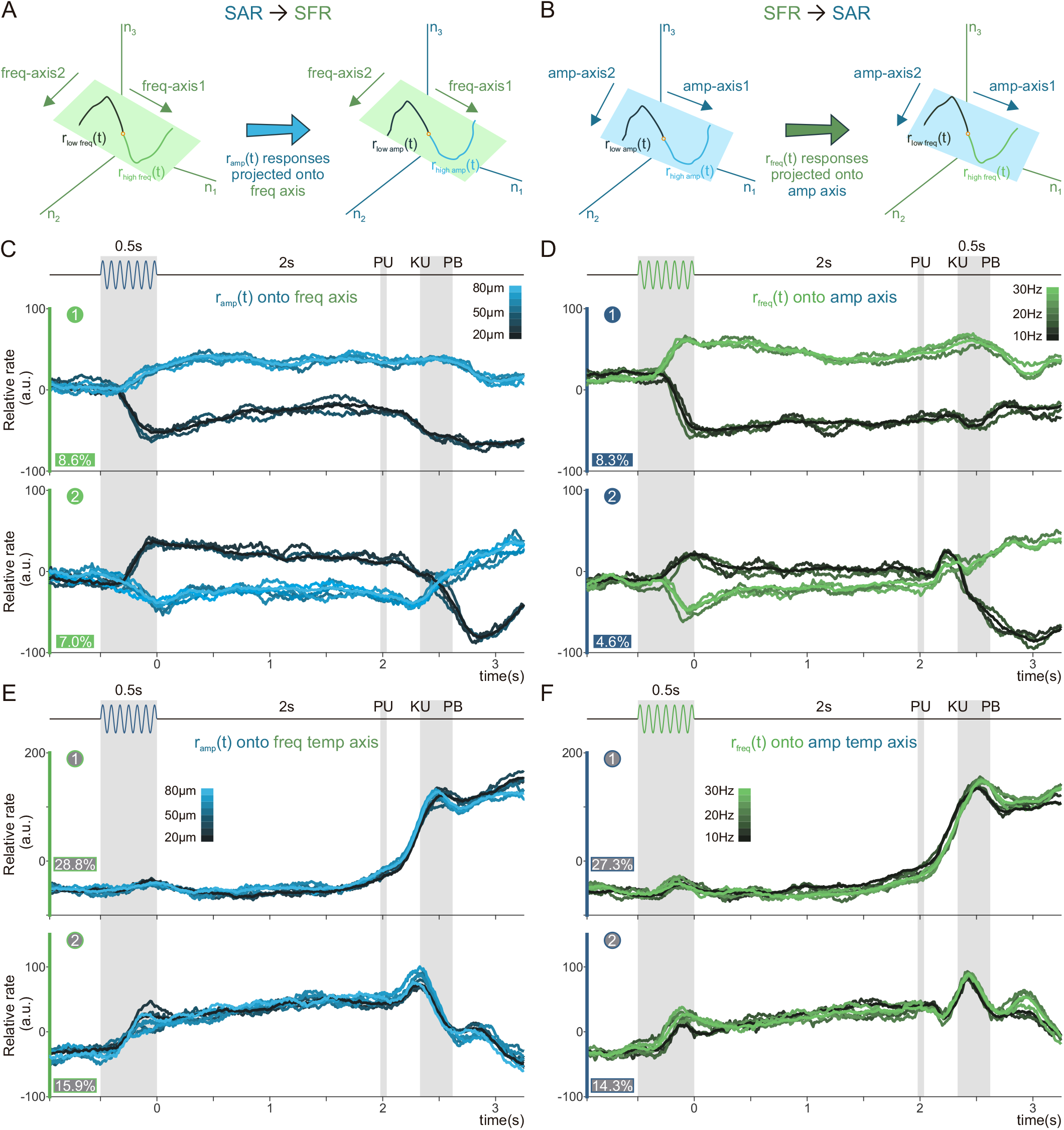
Abstract Temporal and Categorical Coding Population Dynamics. (A, B) Schematic depicting the projection of the short amplitude range (SAR) activity over the short frequency range (SFR) decoding axes (A), and vice versa (B) (n=158). (C, E) Projection of classes from the SAR population over the first 2 SFR freq-axes (C), and the first two SFR temporal-axes (E). Axes computed with the SFR dynamic. The numbers in green circles, as well as the color of the y-axis, mark the source of the decoding axes. Explained variance of the SAR dynamic for each SFR freq-dPC are included below. (D, F) Projection of classes from the SFR population over the first 2 stimuli SAR ampaxes (D), and the first two SAR temporal-axes (F). Axes computed with the SAR dynamics. The numbers in blue circles, as well as the color of the y-axis, mark the source of the decoding axes. Explained variance of the SFR dynamic for each SAR amp-dPC are included below.

### A Continuum of Single Neuron Responses Generates the Population Dynamics

To further characterize the DPC responses, we asked whether the different population dynamics are generated from separate groups of neurons. In other words, is it possible to divide the neural responses into clusters associated with the population responses observed in Fig. 5 and 6? To tackle this question, we paired the neuronal weight distributions to create scatterplots for 3 different pairs of decoding axes: stim-dPC-1 and stim-dPC-2 (Fig. S5A, left), stim-dPC-1 and temp-dPC-1 (Fig. S5A, middle), and temp-dPC-1 and temp-dPC-2 (Fig. S5A, right). In all three cases, the 18 example neurons (Fig. S5B, right and bottom) are distributed in a seemingly random manner, although there are some intuitive differences. For example, neuron units #182 and #007 are relatively closer in the plane created with the stim-dPCs (left), than the one created with the temp-dPCs (right). Intuitively, the response pattern to the varying stimuli is similar for these two example neurons, although they may have two completely different temporal signals supporting their categorical response. This initial analysis supports what other studies have found in other tasks (22, 24, 35): that the population of neurons recorded in DPC appears along a continuum of mixed temporal and stimulus dynamics.

To corroborate these results with a different approach, we used a non-linear dimensionality reduction technique known as Uniform Manifold Approximation and Projection (UMAP) (36). This method was recently applied to identify grid cells in the entorhinal cortex and reveal the toroidal geometry underlying their responses (37). Using this technique, we projected the concatenated firing rate from all stimuli related to the SFR set, into 2-dimensional space, providing us with a visual representation of the data that could potentially detect functional clusters of neuronal activity (Fig. 7A). Under the assumption that such clusters could exist, we first estimated the probability density functions associated to each UMAP dimension (Fig. S5B, above and to the right of the UMAP plane). These nearly unimodal distributions suggest that there is only a single, centralized cluster in the UMAP plane.

**Figure 7.**
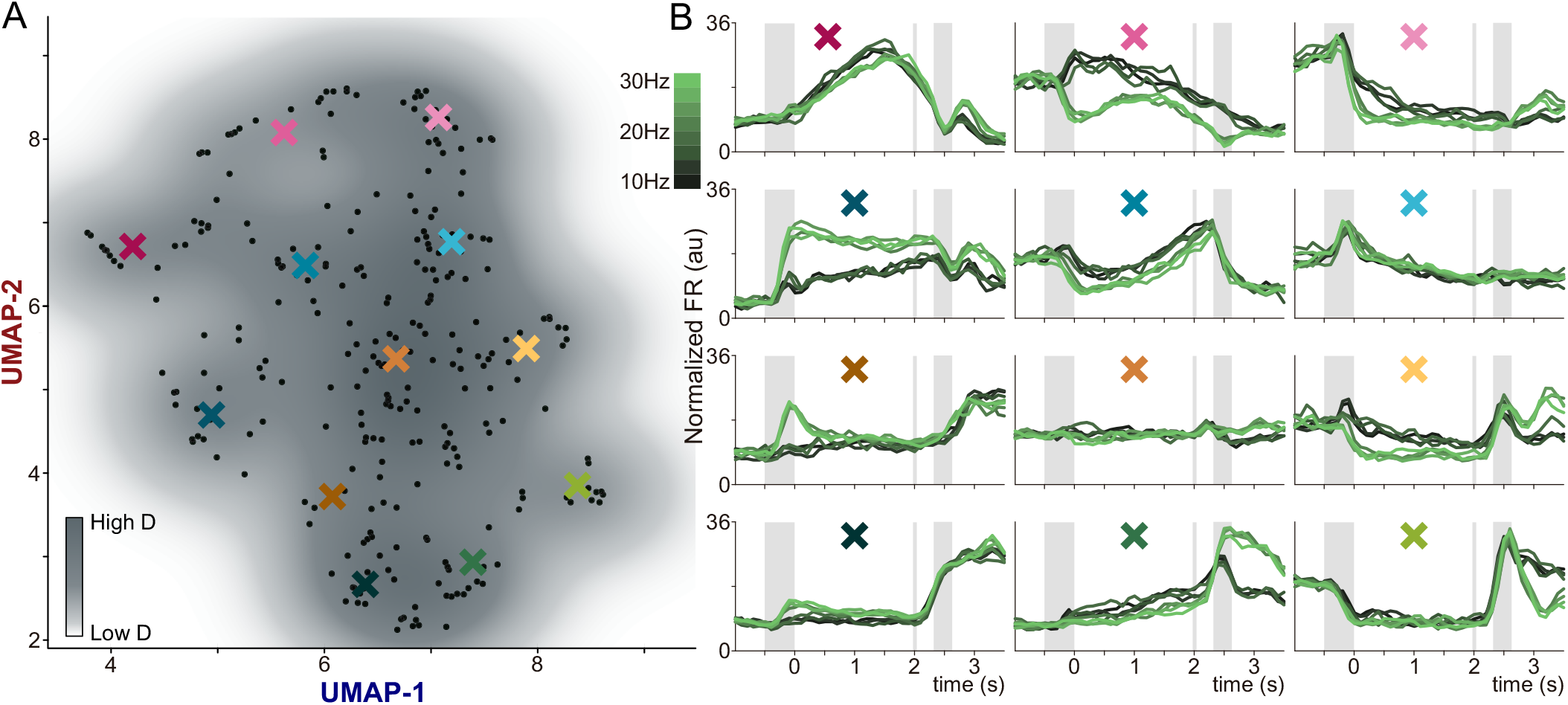
Continuum Interlinked Categorical and Temporal Neural Responses. (A) Uniform Manifold Approximation and Projection (UMAP) plane with a density contour plot. Each black point represents a single neuron. The colored X marks are the centers of the double-Gaussian (σ=0.4) weighted averages presented in the panels in (B). (B) Each sub-panel presents the weighted average of neuronal activity and is presented in back-transformed arbitrary units that roughly correspond to the firing rate values (Hz).

To further address this result, we calculated the peak clustering index (38) (see Methods) of each point and found two outliers (inset in Fig. S5B). When graphing these density peaks (square symbols in Fig. S5), it is possible to note that they do not separate different groups, suggesting that they belong to the same cluster. Moreover, to visualize this single, central cluster, we underlaid a contour density plot onto the neurons plotted across the UMAP plane (Fig. 7A), and observed that our cluster is indeed a single, continuous entity. We sought to characterize how groups of neurons vary across the UMAP plane, so we calculated a double-Gaussian (σ = 0.4) weighted-average of the activity of all neighboring neurons at certain points (X-marks) throughout the UMAP plane (Fig. 7B). Observing the results, we can see that categorical dynamics occur to varying degrees across many of the averages, along with very clear portions of temporal signals. Although these clearly marked and separated periods of categorical versus temporal dynamics are only emerging due to the averaging of several neurons, it provides a unique opportunity to observe how integrally mixed these two regimes of dynamics are. Notably, the inter-trial period demonstrates both categorical and temporal dynamics that have been preserved through the averaging procedure, suggesting the relevance of this DPC signal during this task. From this result, we can conclude that the temporal and categorical dynamics are inherently interrelated, making it impossible to isolate separable functional groups. Ergo, even if the population signals were obtained with orthogonal readouts, they were constructed from the same continuum of neuronal substrates.

## DISCUSSION

In this work, we sought to understand whether the DPC contributes to the elaboration of a generalized, abstract categorical decision code. Two trained monkeys performed the VCT, where vibrotactile stimuli had to be categorized as either “high” or “low”. Two features of the experimental design dictate the core questions of our research: 1) stimuli could vary in either of two different physical attributes: frequency and amplitude; and 2) values for these attributes varied across two superimposed ranges, so that around half of the stimuli switched categories according to context. These two features were developed to test for abstractions in the neural code, and they were motivated by the following type of questions: does DPC generalize its response across both the two ranges and the two physical attributes? This would be the most abstract code possible for this task. Analyzing the neuronal activity of DPC at the single and populational level we found that regardless of attribute or range, the abstract decision is coded by the network employing a unique neuronal population —a common neural substrate— from which an abstract code emerges to represent both categories. From a computational perspective this phenomenon is highly flexible and efficient, given that the same underlying population of neurons arrives at an appropriate decision independently of context. Although the dynamics observed in DPC are not enough to resolve the task by themselves, since they only represent the emerging final decision, the results shown above add to evidence that DPC is not only limited to the planning of physical movement. Importantly, this decision is also coded during the inter-trial period, suggesting that this signal is relevant for future trials. Our results suggest that it is also involved in the construction and reevaluation of a complex and abstract categorical decision code. The notions of a singular abstract code and of a common neural substrate for it are supported by the evidence from d-PCA (24) and UMAP (36). After studying the information carried by single units, these two population approaches were used to examine how this information is distributed across DPC neurons. First, we applied the dimensionality reduction of d-PCA to find the most prominent signals in our population. The foremost finding delivered by d-PCA is that, for all four combinations of range and physical attribute studied, the emergent population signals are the same. This is true for the categorical coding signals, as well as for their underpinning temporal dynamics. This might most clearly be observed when we projected the neural responses to one attribute over the components extracted while the other was presented; doing this swap made no difference to the signals. The implications of this observation are twofold: when we consider that these population signals are obtained by combining all neuronal responses according to a set of weights, on the one hand, this makes evident that the population signals in each context converged to the same abstract categorical code; on the other hand, the weight distributions served as a first indication that there were no separate subpopulations. To further test this, UMAP was brought to bear on the full dynamics of the neural population. Density-based clustering on the results of this non-linear algorithm also showed no presence of subpopulations. Moreover, as it was stated in a previous DPC work (27) and further corroborated here, with UMAP and d-PCA approaches, temporal signals constitute the basic infrastructure over which this abstract code unfolds. Thus, it appears that the singular abstract code emerged from the dynamics of a neural substrate common to all contexts.

Nevertheless, where does DPC fit in the process of constructing this decision? And what signals are available to it? Moreover, are there any upstream areas that are intermediate in their abstraction, generalizing for one feature but not the other? And how would areas produce this abstraction? Do they consolidate subpopulations of neurons that each generalize a certain feature? Or do abstract signals emerge from a common neural substrate? To discuss this, we should consider first where in the somatosensory hierarchy DPC is thought to belong and what the areas upstream to it do. In a variety of purported hierarchies of the somatosensory processing network in macaques, DPC sits between the sensory areas and the motor output (11, 39), together with other premotor association cortices (1). Regarding the sensory areas, the secondary somatosensory cortex (S2) would be immediately upstream to DPC (28, 40); while S1, further away, serves as the entrance into the cortex. Previous work using a VCT has demonstrated how faithful the representation of the physical attributes (amplitude and frequency) is in S1 (12). These findings have been confirmed in a wide variety of tasks (4, 11, 28), as well as several animal models (41, 42). On the other hand, recent results from our group found that S2 could be a suitable candidate for the site of transformation from sensory representations to more abstract signals (40), since both types of responses were observed to have similar proportions in this area. From those findings, S2 appears to be an essential driver of sensory abstraction and the main source of inputs available for DPC to construct its decision categorical code. With this in mind, we do not mean to suggest that DPC is the only area which plays a key role in the formation of this code. Decision-related activity is known to be widely distributed across frontal lobe areas (1), and most are probably also recipients of the signals generated by S2 and equivalent areas from other senses. Nevertheless, previous findings make DPC an interesting target for the study of our research questions (4, 11).

Classically, the role of DPC has been limited to preparing physical motor reports (14, 18, 19, 43), mental rehearsal of known physical movements (13, 44), and serving as an aide to the primary motor cortex (M1) (45–47). Importantly, as it is stated in Methods, our experiment was carefully designed to minimize variations in activity related to push buttons movement. Further, more recent work in this area suggests a gating role, blocking unintended movements while transmitting and allowing the execution of intentional movements (16, 23, 48). This gating of movement signals is inherently linked to the animal’s final decision; similarly, recent work showed that the population activity in DPC codify the entire decision process during a temporal pattern discrimination task (4, 22). Furthermore, there was an additional interesting observation found through our population analyses, reminiscent of previous works (20, 23). Although there was decision-related information before and after the movement (during the inter-trial period), dPCA showed that these two decision signals are arranged in an orthogonal fashion. This might indicate that they have different functions. While the activity during working memory maintains the decision information, the activity after movement could be directly related to an evaluation of the decision to ascertain whether it was rewarded (30). This would mean that DPC is involved in a re-emergence of the decision, with no relationship to movement whatsoever; if the decision-related activity in DPC occurs independently from movement, then its role must be far more nuanced than previously proposed. This could be seen as part of a feedback loop that can continually reinforce the somatosensory processing network’s performance based on recent experience.

However, if previous evidence has shown that abstract categorical decision signals emerge as early as S2, what would be the function of DPC’s decision signal? The answer may lie in the temporal dynamics of the signal. If the frontal lobe does not directly produce these abstract signals, it might be more concerned with a) providing the dynamical infrastructure to sustain abstract information persistently in working memory, b) associating it with relevant contextual information, c) broadcasting the decision in order to make the motor execution, and d) distributing feedback about the outcome afterwards. In our results, points a) and d) might be the most evident; one during the delay between stimulus presentation and decision report, and the other during the inter-trial period. In any case, we would like to highlight a) as a proximate computational answer to the question of DPC’s function: it would seem that this area is not computing the category — that might happen in S2— but rather developing it into an abstract, persistent signal.

In brief, the results presented here show that a common code is employed to maintain the categorization of differing physical attributes each presented in two superimposed ranges. Here we would like to mention that this mechanism for categorical coding, which we encountered is similar to those that have been described for other, different tasks (49). In addition, we have demonstrated through dimensionality reduction techniques, that this singular categorical code emerges from the entire DPC’s neuronal population and not from isolated functional groups. We think that this could be favored thanks to the broad range of heterogeneous responses —a mixture between categorical and temporal dynamics— that single units tend to present (21), which is also known to improve neural network performance (50). Our current work highlights the ability of frontal lobe neurons to adapt strategies to a variety of task conditions. This adaptability can be further studied by testing whether the categorical representation of duration intervals is represented in the same way. Furthermore, although we have observed purely abstracted signals in the DPC activity, the origin of these signals remains to be verified under the structure of the VCT. It is also important to consider whether the working memory activity in DPC could be more nuanced in a task scheme that requires sensory information to arrive at the final decision. Finally, it would be greatly informative to study other kinds of data, such as local field potentials (LFPs), to gain further understanding of how the network utilizes all possible strategies to coordinate categorical decision-related responses.

## ACKNOWLEDGMENTS

We thank Hector Diaz for his technical assistance. This work was supported by grants PAPIIT-IN205022 (to R.R.-P.) and PAPIIT-IG200521 (to V.d.L.) from the Dirección de Asuntos del Personal Académico de la Universidad Nacional Autónoma de México and CONACYT-319347 (to R.R.-P.), CONACYT-319212 (to V.d.L.) and CB2014-20140892 (to R.R.) from Consejo Nacional de Ciencia y Tecnología. G.D.-d.L. (fellowship CONACYT-964544), S.Pa. and J.Z. are doctoral students from Programa de Doctorado en Ciencias Biomédicas, UNAM. L.B. is a postdoctoral student (Postdoctoral fellowship CONACYT-838783).

## MATERIALS AND METHODS

### Vibrotactile Categorization Task

This task has been previously described in (12). Two monkeys *(Macaca mulatta)* were trained to label the intensity of different physical attributes (frequency or amplitude) of a vibrotactile stimulus as “high” or “low”. The monkeys were presented with blocks of trials, known as ‘‘sets’’, using predetermined groups of stimuli. Before each set, the monkeys were presented with the lowest and highest stimuli of the set to indicate the physical attribute to be categorized, as well as the range of categorization. The monkeys were trained in the categorization task using two different ranges for each attribute: this means that a total of 4 sets were used throughout the experiment. Neither the range nor the variable attribute changed throughout a set. Monkeys were seated in a primate chair with their right arm, hand, and fingers comfortably restrained. The left hand operated an immovable key (elbow at ~90°), as well as two push-buttons, separated 11 mm from their edges, located directly in front of the animals, 25 cm away from the shoulder, and at eye level. Stimuli were applied to a distal segment of a single digit of the restrained right hand via a computer-controlled stimulator probe (2mm round tip, BME Systems, Baltimore, MD).

Vibrotactile stimuli were composed of short mechanical pulses, each consisting of a 20ms single-cycle sinusoid. All stimuli lasted 500 ms and varied in either frequency (number of pulses) or amplitude (depth of indentation on the skin). Time was always aligned to the first stimulus offset (0s corresponds to the end of the stimulation period). Each trial began with the probe down event (“PD”), when the probe descends and indents the skin with a depth of 500 μm (Fig. 1). This was followed by a waiting period of random length (1.5-3s), to prevent the monkeys from predicting the arrival stimuli. Afterwards, the 500ms of stimulation were delivered. Then, monkeys were presented with a fixed 2s delay. The end of this delay was marked by the probe up event (“PU”), which served as the “GO” cue for the monkeys to release the immovable key (key up, “KU”) and report their decision using one of the two push-buttons (push-button, “PB”). The rewards for correct decisions were drops of fruit juice. Animals were handled in accordance with standards of the National Institutes of Health and Society for Neuroscience. All protocols were approved by the Institutional Animal Care and Use Committee of the Instituto de Fisiología Celular of the National Autonomous University of Mexico (UNAM).

Because we were interested in finding cortical activity related to the decision, particularly in DPC, it was crucial to minimize or eliminate modulatory effects arising from the well-known dependence of arm movement direction (1, 51) or on parameters that covary with it. The setup was thus arranged to filter out directionally tuned responses. The distance between the push buttons was 3.5 cm, and these were 18 cm away from the immovable key. Directional neurons fire at frequencies that range between ~5 and 25 spikes per s (1, 51), corresponding to their antipreferred and preferred directions, respectively. Therefore, on average, directional neurons modulate their firing rates by ~25 spikes per s when movement direction changes by 180°. The expected effect on an 11° change in direction is thus on the order of one spike per s. Under these conditions, some activity related to arm motion may be expected but should be practically identical for the two arm movements.

#### Set Design

It is important to explain some important aspects of the sets’ design. First, the two different stimulus ranges tested with each attribute were of “short” and “long” ranges. Thus the 4 sets used were: short frequency range (SFR), long frequency range (LFR), short amplitude range (SAR) and long amplitude range (LAR). The SFR and SAR (short sets) had 12 stimuli, with frequencies going from 10 to 30 Hz and amplitudes from 20 to 80 μm; the LFR and LAR (long sets) had 10 stimuli, with ranges of 14-78 Hz and 42-138 μm. In this way, the top half of the short sets roughly correspond to the bottom half of the long sets. This means that those stimuli e.g. 30 Hz, would be considered high in the short set but low in the long one, forcing the monkeys to change their criterion. Further, the monkeys could not construct a criterion based on absolute differences, since the long sets had larger intensity differences between stimuli than the short sets. Second, in all sets the threshold stimulus, which had the intensity value at the middle of the range and thus separated the categories, was always presented twice as many times. Half of the time the rewarded decision was to label it “high” and the other half it was “low”. This ensured that the threshold stimuli remained perceptually ambiguous during each set. Importantly, it also allows for the study of the two different perceptual judgments with exactly the same stimuli.

#### Training

Both animals were first trained with the short sets (SFR and SAR) until they reached a performance of 80%. Once this level of mastery was maintained for the short sets, the long sets were introduced (LFR and LAR). Then, when animals sustained this reasonable level of performance across all 4 sets, neuronal activity recording began. Across each day of recording the order of our 4 sets (the combinations of attributes and ranges) varied randomly.

##### Recordings

Neuronal recordings were obtained with an array of seven independent, movable microelectrodes (2-3 MΩ) inserted into the dorsal premotor cortex (DPC), either contralateral (left hemisphere) or ipsilateral (right hemisphere) to the stimulated hand. We collected neuronal data in blocks using the predetermined sets, with a minimum of 5 trials presented per stimulus intensity. Neurons that maintained similar response patterns across recordings of different sets were said to be the same only if the recordings occurred along the same electrode and the electrode’s position was not moved. All recordings in DPC were made in the arm region of F2 (1, 4). This region is in front of M1 (F1), lateral to the central dimple, posterior to F7 and the genu of the arcuate sulcus (52). Recording sites changed from session to session and the locations of the penetrations were used to construct surface maps of all of the penetrations in each cortical area. This was done by marking the edges of the small chamber (7 mm in diameter) placed above each cortical area. We recorded a total of 275 neurons for SFR, 232 neurons for SAR, 84 for LFR and 59 LAR.

##### Data Analysis

###### Firing Rate

For each neuron, we calculated a time-dependent firing rate for each trial using overlapping windows of 200ms separated by 10ms steps. To produce the class-firing rate curves (Fig. 2), we averaged the rates of correct trials from each stimulus intensity.

###### AUROC

For each neuron, only correct trials were taken into consideration, and all trials were grouped based on their associated category. For threshold stimuli, trials were sorted according to the monkeys’ behavioral response. At each time bin we defined two distributions, one for category low and one for category high, and calculated a running-integral to produce the Receiver-Operating Characteristic (ROC) curve (2, 4). Integrating the ROC curve yields values that range between 0 and 1: a value of 0.5 indicates that the two categorical distributions were perfectly overlapped; less than 0.5 denotes a preference for greater responses to category low classes; more than 0.5 denotes greater responses to high classes.

###### Mutual Information

For each neuron, we calculated two different types of mutual information for each time window: stimulus information (I_STIM_) and category information (I_CAT_). To compute I_CAT_, we considered only correct trials and divided these into two groups, based on their associated category. Again, we sorted threshold stimuli based on the monkeys’ behavioral report. With this, we produced two distributions of firing rate values for each time bin: P(r|Low), associated with category low, and P(r|High) with category high. We also computed a third distribution, P(r), which quantified the probability of any given response.

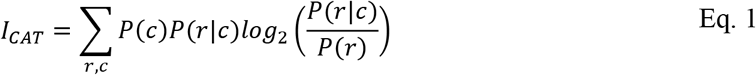

To compute I_STIM_, we grouped the trials according to the stimulus value. We computed one firing rate distribution for each group, as well as a sixth distribution for all responses, P(r).

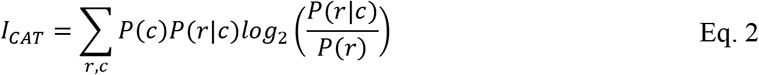

###### Permutation Test for Significance

To evaluate the significance of AUROC and mutual information values we performed permutation tests with 1000 iterations. For each iteration we shuffled the stimulus labels randomly and then recalculated the metric. Next, we counted the number of times in which a metric’s shuffled value was greater than the original, and finally divided it by the number of iterations to obtain a p-value. For mutual information we used a cutoff of 0.05 for significance and one of 0.01 for the AUROC results. For all mutual information values computed across this work, a correction for sampling bias was applied (53). Further, for its significance test we performed a correction for multiple comparisons with a clustering method (54). This was done by maintaining only the set of significance-connected time bins with a size bigger than a threshold.

###### Population Analysis

The responses found within the population of DPC neurons during the categorization task were heterogeneous and demonstrated association to several different task parameters (22). To better elucidate the dynamics at play, we analyzed the neuronal population signals. First, we averaged the firing rate of all correct trials for each of the different stimulus intensities within a single set. Using these class-averaged firing rate curves, we constructed pseudo-simultaneous population responses, in such a manner that the response at every time to each stimulus intensity was represented by a separate N-dimensional vector *<u>r</u>*(*t,s*), also known as a population vector. Each component of the N-dimensional vector represents the activity of one neuron i.e. N refers to the number of neurons in the population. In order to study the categorical representations in the population, we created the population vectors *<u>r</u>*(*t, Low*) and *<u>r</u>*(*t, High*) from the average of hit trials from each category. Finally, we created a population vector that averaged all hit trials (*<u>r</u>*(*t*)). All population vectors represent a single point in time, so we grouped them in a time dependent manner to create a population matrix that summarized all the neurons’ activity associated with either a single stimulus intensity or a single category.

###### Demixed Principal Component Analysis (dPCA)

The details regarding the underlying algorithm, as well as related mathematical proofs, were laid out in (24). dPCA is a dimensionality reduction technique that takes diverse dependencies into account, allowing us to study the population coding of different task parameters and the pure temporal signals. The method has a supervised and an unsupervised portion. In general terms, the supervised portion of the algorithm is related to the selection of relevant task parameters. These task parameters are used to calculate marginalized covariance matrices associated to the individual parameters, as well as to the interaction between sets of those same parameters. Once the marginalized covariance matrices are obtained, the algorithm proceeds to execute an analysis similar to that of PCA, identifying dimensions that explain the greatest amount of variance associated with each of the parameters selected. This portion of the algorithm is unsupervised. Due to the dependence of the categories on the stimulus intensities, it was not possible to separate the activity associated to the stimuli from the activity associated to the categories. For this reason, we performed two separate variants of dPCA: stimulus-dPCA and category-dPCA. For the former, we marginalized the total population activity (<u>*X*</u>) with respect to time 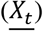 and with respect to stimulus identity 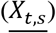. For the latter, we marginalized the population activity with respect to time 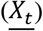 and then to category identity 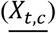. We calculated the marginalization averages as follows:

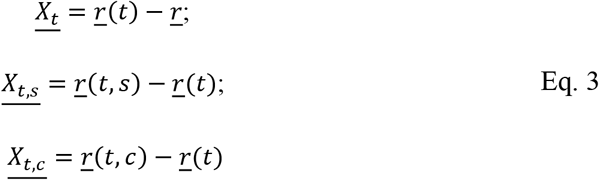

The resultant matrix, 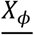, denotes the marginalized data matrices associated to one of the two types of dPCA, such that *ϕ*∈ (*t*, {*t, c*}} in the case of category-dPCA, and *ϕ*∈ (*t*, {*t, s*}} in the case of stimulus-dPCA. In the equations above, *r* is a population vector with the average firing rate values taken over the entire task period (−1 to 3.2s), *<u>r</u>*(*t*) is the population vector with the average firing rates of all hit trials at a single time point, *<u>r</u>*(*t, c*) is the population vector with the average firing rates per category at a single time point, and *<u>r</u>*(*t, s*) is the population vector with the average firing rates per stimulus identity at a single time point. Each of these population vectors is N-dimensional, with a length equal to the total number of recorded neurons. Further, *r^i^*(*t*) represents the firing rate average across all trials at time t for neuron i. Once the marginalized matrix was obtained, the algorithm calculated the decoder (dimensionality reduction) and encoder (recovery transformation) matrices for each 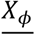, by utilizing reduced-rank regression to minimize the term:

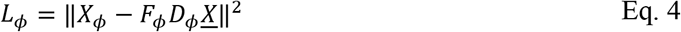

In the equation above, *<u>X</u>* is the centered whole population data matrix, such that the average activity of each neuron is 0. The analytic solution can be found using singular value decomposition (24). *F_ϕ_,D_ϕ_* represent decoding and encoding pair for a given variable, *ϕ*. With respect to our usage, *φ* represents the individual task parameters. Decoder and encoder axes constitute the dimensionality reduction step that reduces the data into the fewest, most representative components, associated to the individual task parameters. In turn, the original neural activity can be reconstructed through linear combinations of these components, just as in PCA. The components of *D_ϕ_* can be ordered in terms of the amount of explained variance. The first component, which explains the most variance, is known as the 1^st^ demixed principal component (1^st^ dPC). Since overfitting is an issue to be considered when employing dPCA, a regularization term was introduced, and cross-validation was performed to choose the most appropriate regularization parameter.

To obtain Figure 5, we projected the N-dimensional data for a given class (*<u>r</u>*(*t, s*)) onto the most prominent decoding axis (k) of a given variable *ϕ*, using stimulus-dPCA (Fig. 5A-B,C,E,G-H). These projections were computed using the following equation:

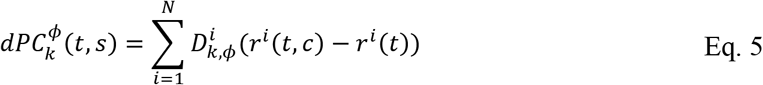

We also produced distributions of the individual neuronal weights that compose each of the stimulus parameter decoding axes (Fig. 5D,F). These same distributions are combined to create planes in Fig. S5A, showing that no observable clustering can be identified using the neuronal weights. This provides evidence that dPCA is not capable of separating functional clusters from the data provided.

For Figure 6, we calculated the decoding axes from a fixed population recorded for both short sets, while maintaining a consistent number of trials between the two compared populations. As a result, we were able to do “cross-projections”: projecting the population vectors from the frequency set onto the decoding axes of the amplitude one (A,C,E), and vice versa (B,D,F). The same procedure was used to obtain Fig. S4, although with the long sets instead of the short ones.

###### Uniform Manifold Approximation and Projection (UMAP)

This algorithm is a non-linear dimensionality reduction technique, based on: (1) approximating the manifold on which we assume the data was sampled from and (2) projecting this manifold into a low dimensional space while maintaining its local and global topological properties (36). For this, we concatenated all the *<u>r</u>*(*t, s*) vectors of the 12 stimuli intensities corresponding to the SFR set horizontally for each neuron. The firing rate was calculated using a window of 200ms and steps of 100ms, yielding a total of 51 time-bins per stimuli intensity (class), and 612 dimensions once all vectors were concatenated. As a result, the size of the matrix input into UMAP was 275×612 for the SFR set, where the 275 neurons were taken as observations, and the 612 time-point dimensions were reduced to two representative dimensions. We then project the inferred manifold into the new 2dimensional space in order to determine whether a non-linear dimensionality reduction technique could isolate neural responses with the characteristic population dynamics that we have identified in Fig. 5 and 6. In Fig. 7A, we included a density contour plot calculated with a Gaussian kernel (σ=0.5). For Fig. 7B, we calculated a double-Gaussian (σ = 0.5) weighted-average centered on each of the colored X marks to visualize how the dynamics vary across the UMAP plane: neurons were weighted based on the Gaussian PDF and then divided by the total sum of the weights.

###### Clustering Index

To identify possible groups of neurons with similar coding or dynamic properties we performed a density cluster analysis (38). In brief, the procedure consists of determining the local density of every point, in our case with a Gaussian kernel (σ = 0.5), and its distance to another point with higher density. The peak index is the product of the density and the distance for each point. The points are then sorted by their peak clustering value in descending order (Fig. S5B inset). Cluster centers, characterized by high local density and distance to other peaks should appear as outliers.

---

**Figure S1.**
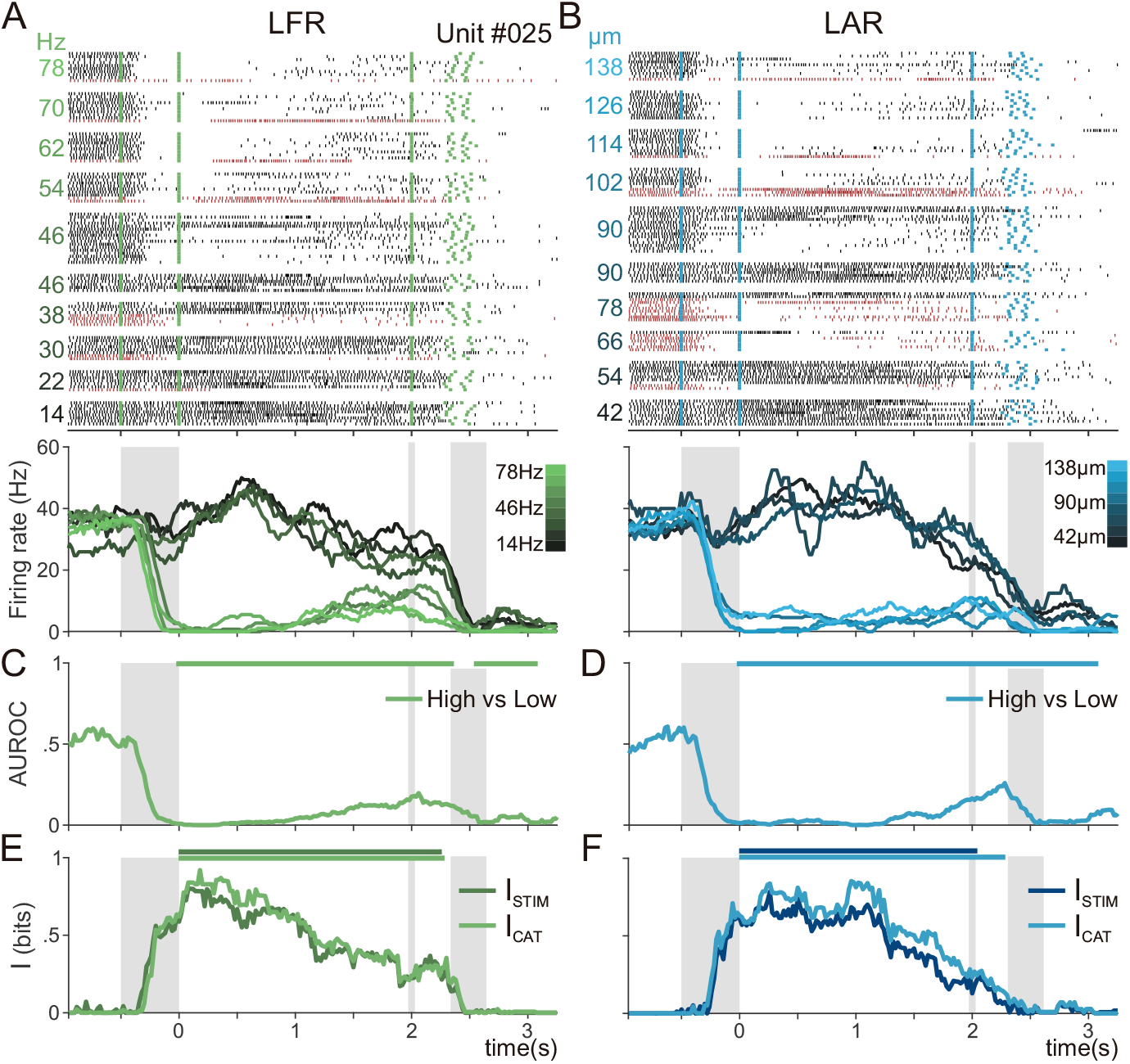
A single neuron’s activity during frequency and amplitude long-range categorization. (A, B) Raster plots of a single neuron recorded in the frequency (LFR; A, C, E) and amplitude (LAR; B, D, F) long-range sets. Green (A) and blue (B) tick marks indicate psychophysical events. Trials are grouped by class (stimulus intensity), with intensity values marked on the left side. Threshold classes are sorted based on subject’s psychophysical report. Firing rate averages for all hit trials associated with each of the 8 visualized classes are shown below. Firing rate curves range from dark grey for low category stimuli to bright green (A, frequency) or bright blue (B, amplitude) for high stimulus category. (C, D) Differential AUROC (High vs Low) values taken between the firing rate distributions of correct trials for category low vs those for category high, in a time-dependent manner. Significant windows, based on permutation tests and correcting for multiple comparisons, are marked with a circle above the curve (p<0.01). (E, F) Category mutual information (I_CAT_, light green and blue) and stimulus mutual information (I_STIM_, dark green and blue) curves are represented in a time-dependent manner (p<0.05).

**Figure S2.**
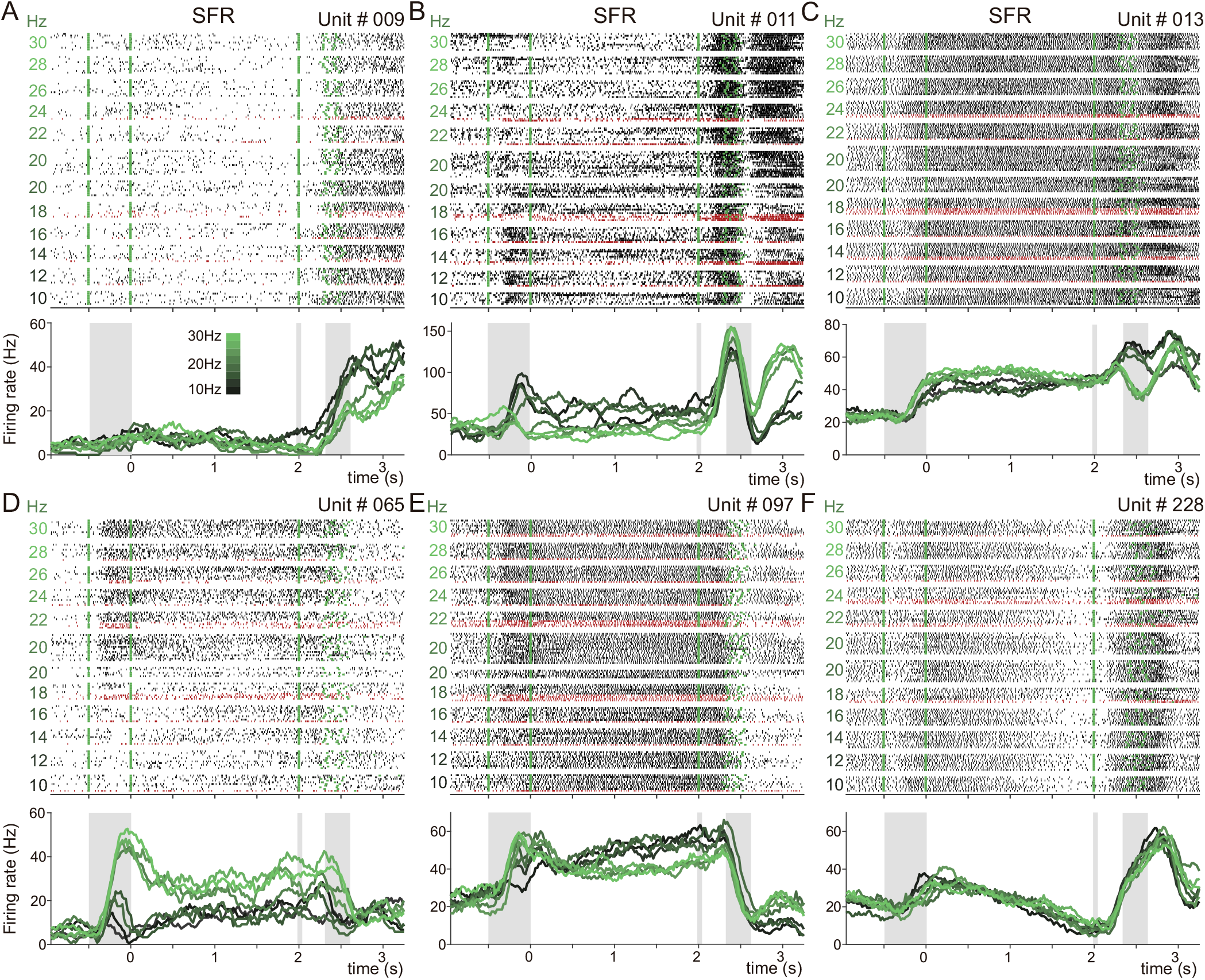
Heterogenous single neuron responses recorded during frequency categorization. (A-F) Raster plots of 6 different neurons recorded in the short frequency range set (SFR). Same format as in Fig. 2 and S1. (A) A neuron with categorical coding that only emerges during movement and lasts into the inter-trial period. (B) A neuron with early categorical dynamics, but no persistent maintenance, which in turn presents the opposite categorical preference during the inter-trial period. (C) A neuron with a nearly pure temporal signal that exhibits categorical responses during movement. (D) A neuron with a preference for stimulus category high that maintains a single representation throughout the delay period. (E) A neuron with categorical dynamics that emerge during the delay are maintained until just after the delay period finishes, and then a slight categorical coding during the intertrial. (F) A neuron with pure temporal signals and no categorical dynamics.

**Figure S3.**
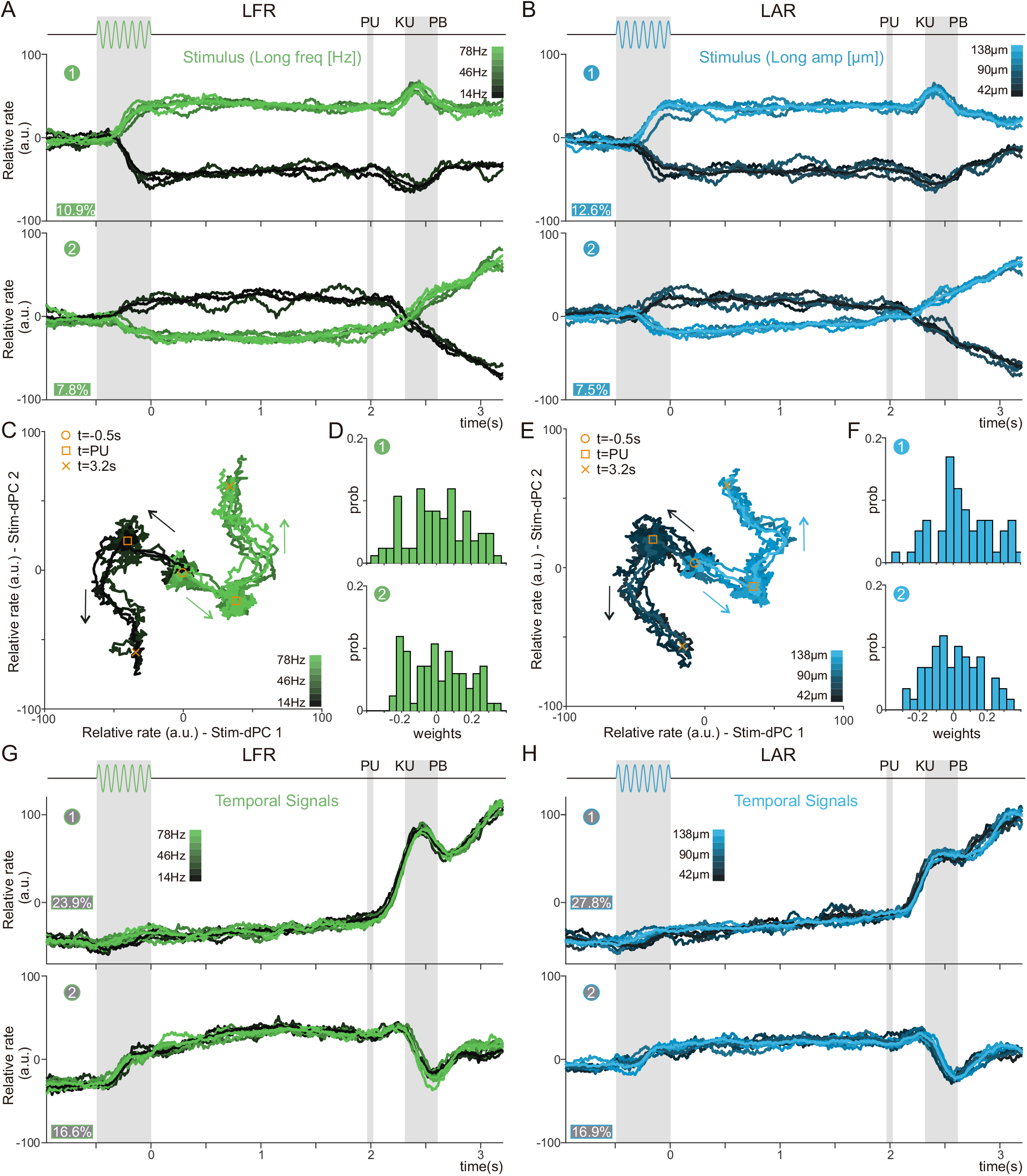
Stimulus and Temporal Population Dynamics during Long-Range Frequency and Amplitude sets. (A, B) Projections of class average over the first and second decoding stimulus axes of the long-range population for frequency (A, LFR, n=84) and amplitude (B, LAR, n=59). Classes range from dark (lowest stimulus) to light (highest stimulus). The ordinal number of each dPC is shown in a circle; explained variances are shown as percentages. (C, E) Phase diagram of the same classes projected across the first 2 decoding axes for frequency (C) and amplitude (E). The yellow circle marks t=-0.5s (SO), the yellow square marks the end of the delay (PU), and the yellow x marks t=3.2s (intertrial period). (D, F) Distributions of the neuronal weights for the first 2 decoding axes of long frequency (LFR; D) and amplitude range (LAR; F). (G, H) The first two temporal population signals for the LFR (G) and LAR (H) populations.

**Figure S4.**
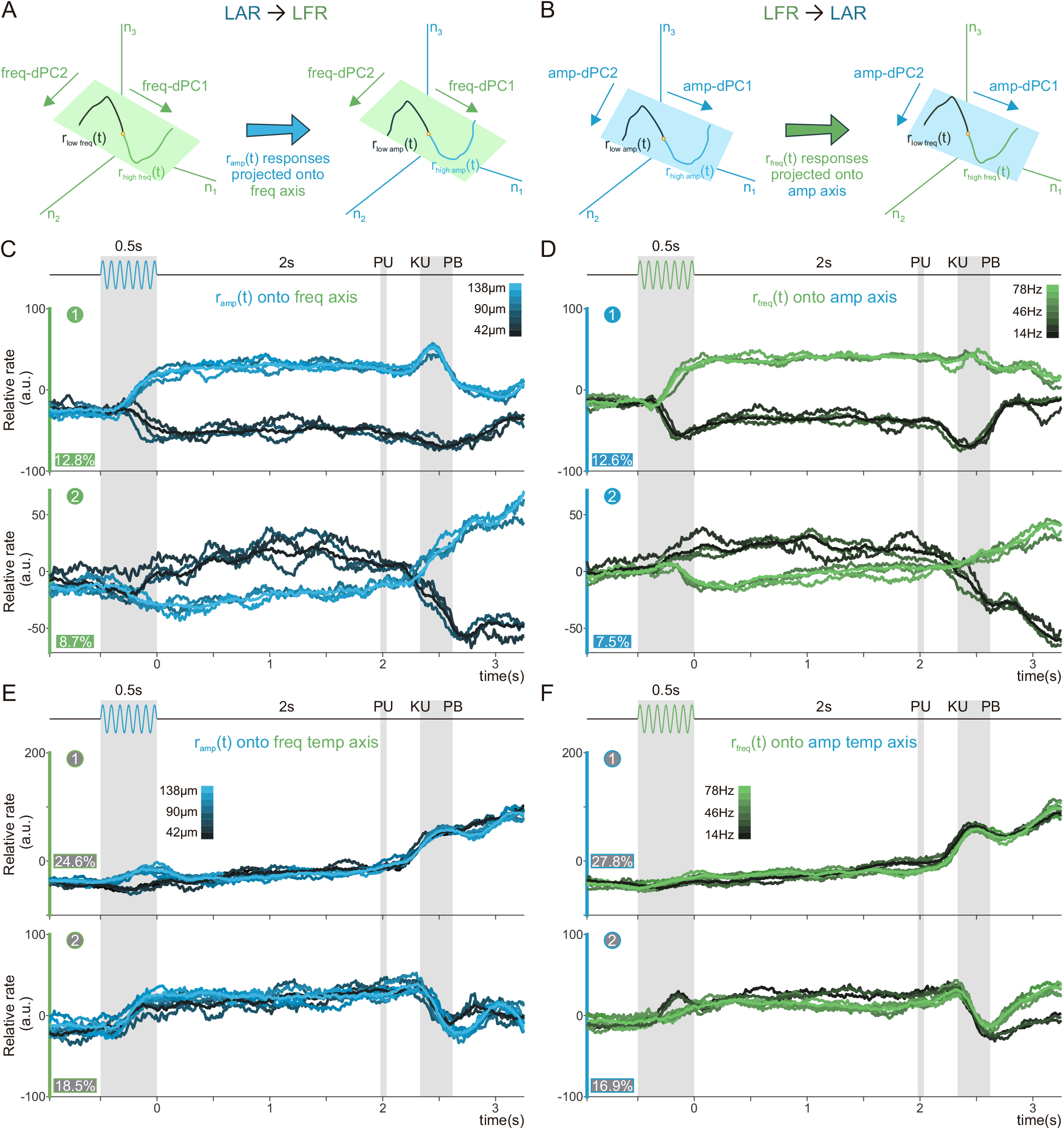
Abstract Temporal and Categorical Coding Population Dynamics during Long-Range Sets. (A, B) Schematic depicting the projection of the long amplitude range (LAR) activity over the long frequency range (LFR) decoding axes (A), and vice versa (B) (n=57). (C, E) Projection of classes from the LAR population over the first 2 stimuli LFR freq-axes (C), and the first two LFR temporal-axes (E). Axes computed with the LFR dynamics. The numbers in green circles, as well as the color of the y-axis, mark the source of the decoding axes. Explained variance of the LAR dynamic for each LFR freq-dPC are included below. (D, F) Projection of classes from the LFR population over the first 2 stimulus LAR amp-axes (D), and the first two LAR temporal axes (F). Axes computed with the LAR dynamic. The numbers in blue circles, as well as the color of the y-axis, mark the source of the decoding axes. Explained variance of the LFR dynamic for each LAR amp-dPC are included below.

**Figure S5.**
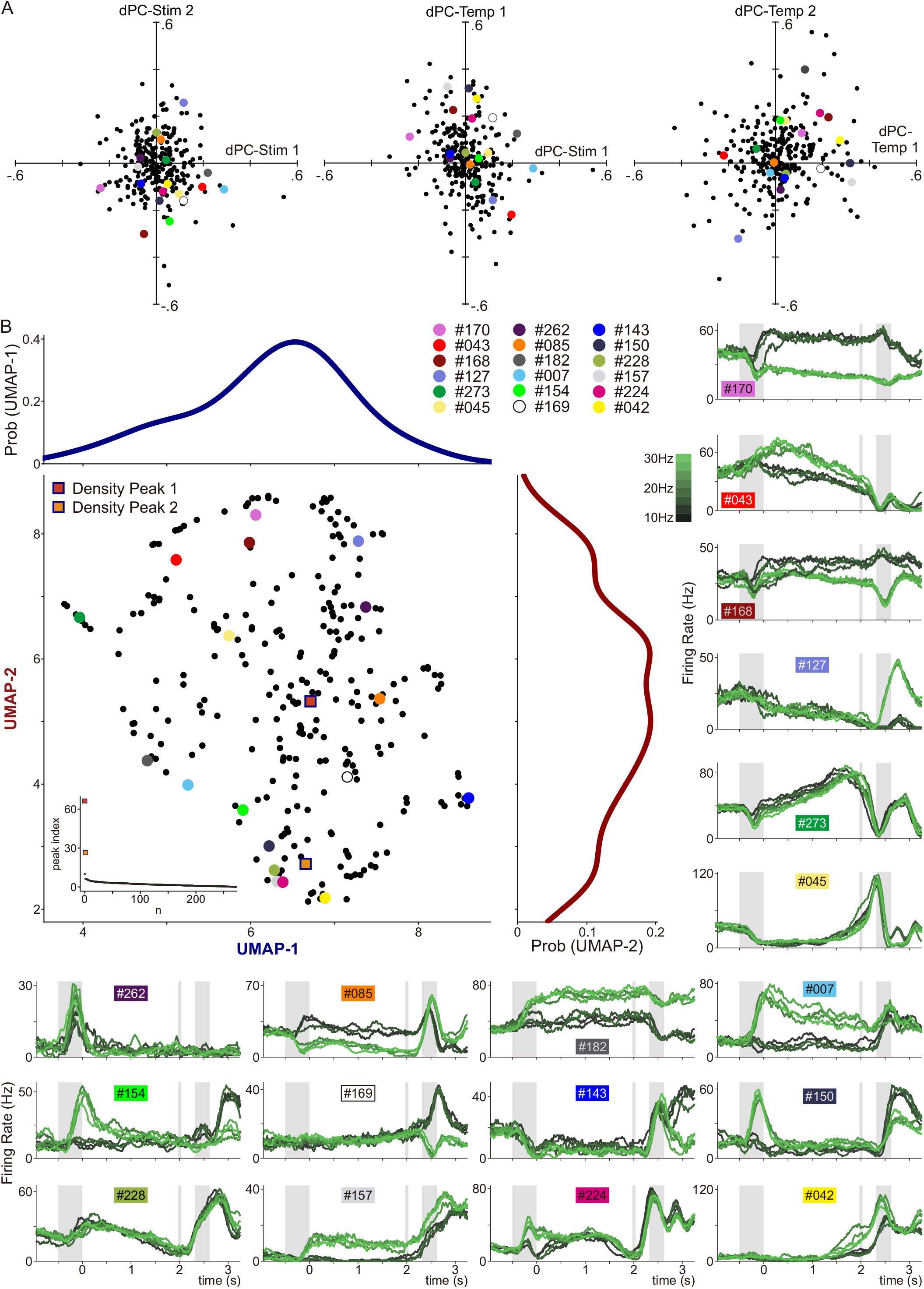
Dimensionality Reduction does not Separate Functional Groups. (A) Scatter plot of the neuronal weights associated with two different dPCA axes during SFR; left: the 1st and 2nd frequency axis (freq-dPC); center: the 1st freq-dPC and the 1st temporal (temp-dPC); right: the 1st and 2nd temporal axis (temp-dPC). Points represent neurons with colored points corresponding to the unit labels that appear in sub-panels of (B). (B) UMAP plane identical to the one presented in Fig. 7A, and corresponding probability density functions for the first (above; dark blue) and second (right; red) UMAP dimensions. Neuron unit colors are presented in the upper-right hand corner of the UMAP plane plot. The inset graph shows the density peak values, sorted in descending order, with two clear density peak outliers (red and orange squares are the density peaks 1 and 2 respectively). Starting with unit #170 in the top right-hand corner, firing rates of 18 heterogenous neurons are displayed in the same format as in Figs. 2, S1 and S2.

